# Monovalent and trivalent VSV-based COVID-19 vaccines elicit potent neutralizing antibodies and immunodominant CD8+ T cells against diverse SARS-CoV-2 variants

**DOI:** 10.1101/2022.07.19.500626

**Authors:** Kate A. Parham, Gyoung Nyoun Kim, Nasrin Saeedian, Marina Ninkov, Connor G. Richer, Yue Li, Kunyu Wu, Rasheduzzaman Rashu, Stephen D. Barr, Eric J. Arts, S.M. Mansour Haeryfar, C. Yong Kang, Ryan M. Troyer

**Author notes:** Corresponding author: Ryan Troyer.

## Abstract

Recombinant vesicular stomatitis virus (rVSV) vaccines expressing Spike proteins of Wuhan, Beta and/or Delta variants of SARS-CoV-2 were generated and tested for induction of antibody and T cell immune responses in mice. rVSV-Wuhan and rVSV-Delta vaccines and a rVSV-Trivalent (mixed rVSV-Wuhan, -Beta, -Delta) vaccine elicited potent neutralizing antibodies (nAbs) against live SARS-CoV-2 Wuhan (USAWA1), Beta (B.1.351), Delta (B.1.617.2) and Omicron (B.1.1.529) viruses. Prime-boost vaccination with rVSV-Beta was less effective in this capacity. Heterologous boosting of rVSV-Wuhan with rVSV-Delta induced strong nAb responses against Delta and Omicron viruses, with rVSV-Trivalent vaccine consistently effective in inducing nAbs against all the SARS-CoV-2 variants tested. All vaccines, including rVSV-Beta, elicited a spike-specific immunodominant CD8^+^ T cell response. Collectively, rVSV vaccines targeting SARS-CoV-2 variants of concern may be considered in the global fight against COVID-19.

## Introduction

The ongoing global coronavirus disease (COVID-19) pandemic caused by severe acute respiratory syndrome coronavirus 2 (SARS-CoV-2) infection has resulted in over 559M cases worldwide and over 6.36M deaths as of July 18, 2022 (1). Infection waves have been driven by viral variants defined by mutations throughout the viral genome with specific mutations in the spike protein that appear to contribute to immune escape in the human population, i.e. SARS-CoV-2 Delta (B.1.617.2), and the current variant of concern Omicron (B.1.1.529). Emergence of new SARS-CoV-2 variants highlights the urgency to develop safe, efficient, and cost-effective vaccines to vaccinate and boost the world’s population, not only to prevent disease, but to impede the emergence of new variants of concern.

Vesicular stomatitis virus (VSV) is a member of the Rhabdoviridae family and has been effectively manipulated to express target antigens on the viral surface, in combination with or replacing the VSV envelope glycoprotein, to generate an immunogenic vaccine vector (2). VSV is a safe vaccine platform in humans and effectively induces immunity in the mucosa and systemically (3). Unlike other viral vector platforms (e.g. adenovirus), VSV does not commonly infect humans, and therefore pre-existing seropositivity is low in the population, minimizing the potential effect of anti-vector immunity which can hamper repeated boosting (4,5). Moreover, VSV can be modified to maintain attenuated replication (6), resulting in prolonged antigen presentation and lower doses per vaccination, decreasing manufacturing requirements and costs when compared to replication incompetent vaccine vectors.

The first FDA approved recombinant VSV (rVSV) vaccine is Ervebo, a live, attenuated rVSV vaccine presenting the envelope glycoprotein of Zaire ebolavirus, which is 100% effective at preventing laboratory-confirmed Ebola virus disease in confirmed contacts of Ebola virus patients (7). Our group and others have utilized the rVSV vaccine platform in the current pandemic for the generation of SARS-CoV-2-specific rVSV vaccines (8–10). These rVSV vaccines utilize spike from the SARS-CoV-2 outbreak founder strain (Wuhan) to induce neutralizing antibodies (nAbs) and spike-specific T cell responses, which greatly reduce virus titers upon infection, result in rapid viral clearance and prevent severe disease in animals following challenge with the Wuhan strain (8–11). BriLife® a replication-competent, Wuhan spike-only rVSV vaccine is in Phase 2 clinical trials for COVID-19, generating immune sera cross-reactive to emerging SARS-CoV-2 variants, including Delta and Omicron (12). These studies show rVSV is an effective platform for the generation of SARS-CoV-2 vaccines against the original SARS-CoV-2 strain. With the emergence of new highly mutated SARS-CoV-2 variants there is a need to generate a broad immune response to vaccination, which could be achieved by developing booster vaccines using variant spike sequences.

Using our previously validated rVSV SARS-CoV-2 Wuhan (rVSV-Wuhan vaccine) prime-boost immunization strategy (8), we generated rVSV vaccines containing full length spike genes from SARS-CoV-2 B.1.351 (rVSV-Beta vaccine) and SARS-CoV-2 B.1.617.2 (rVSV-Delta vaccine). Mice were prime-boost immunized with various combinations of the three monovalent variant vaccines and a trivalent vaccine (rVSV-Trivalent vaccine), containing rVSV-Wuhan vaccine, - Beta vaccine, and -Delta vaccine. Here, we examine the immune response induced in rVSV-vaccinated animals against live SARS-CoV-2 USAWA1 (SARS-CoV-2_Wuhan_USAWA1_), B.1.351 (SARS-CoV-2_Beta_), B.1.617.2 (SARS-CoV-2_Delta_), and the current variant of concern B.1.1.529 (SARS-CoV-2_Omicron_). Monovalent and trivalent immunization regimens elicited nAbs effective against diverse SARS-CoV-2 variants. Further spike-specific CD8+ T cell responses were confirmed with all vaccination regimens. Herein, we show rVSV variant spike vaccines may be an important tool as a booster vaccination in a landscape of emerging SARS-CoV-2 variants.

## Methods

### Cells

Vero E6 cells were obtained from ATCC (ATCC, CRL-1586) and Vero cells obtained from Dr. Louis Flamand, Université Laval. Cells were cultured in complete Dulbecco’s modified Eagle’s medium (DMEM, Gibco) with 10% fetal bovine serum (FBS), 1x Glutamax, and 100 units/ml penicillin/100 μg/ml of streptomycin (Pen/Strep). BHK-21 cells (ATCC, CCL-10) were maintained in DMEM containing 5% FBS, 2 mM L-glutamine, and Pen/Strep.

### Construction of rVSV-SARS-CoV-2 variant vaccines

To construct SARS-CoV-2 vaccines against the Beta and Delta variants, we introduced mutations into the human codon-optimized full-length S (S_F_) protein of SARS-CoV-2 (GenBank: JX869059.2) along with honeybee melittin signal peptide (msp) replacing the S_F_ amino terminus and VSV G protein transmembrane domain and cytoplasmic tail (Gtc) replacing the S_F_ carboxy terminus as previously described (8). The beta variant vaccine (rVSV-Beta vaccine) contains the K417N, E484K, and N501Y mutations in the receptor-binding domain (RBD) of codon-optimized S_F_ (**Figure 1**). The delta variant vaccine (rVSV-Delta vaccine) contains 9 amino acid changes (T95I, G142D, E156G, A222V, L452R, T478K, D614G, P681R, D950N) and two amino acid deletions at F157 and R158, in the codon-optimized S_F_ (**Figure 1**). Mutations were based on predominant circulating spike sequences for each variant (13). The S_F_ genes with mutations were synthetically generated (Genscript USA Inc.) and inserted between the glycoprotein (G) and polymerase (L) genes of VSV (serotype Indiana) at Pme I and Mlu I restriction sites (**Figure 1**). We recovered each rVSV virus by VSV reverse genetics and purified the virus with three consecutive rounds of plaque picking and amplification in BHK-21 cells (8,14).

**Figure 1:**
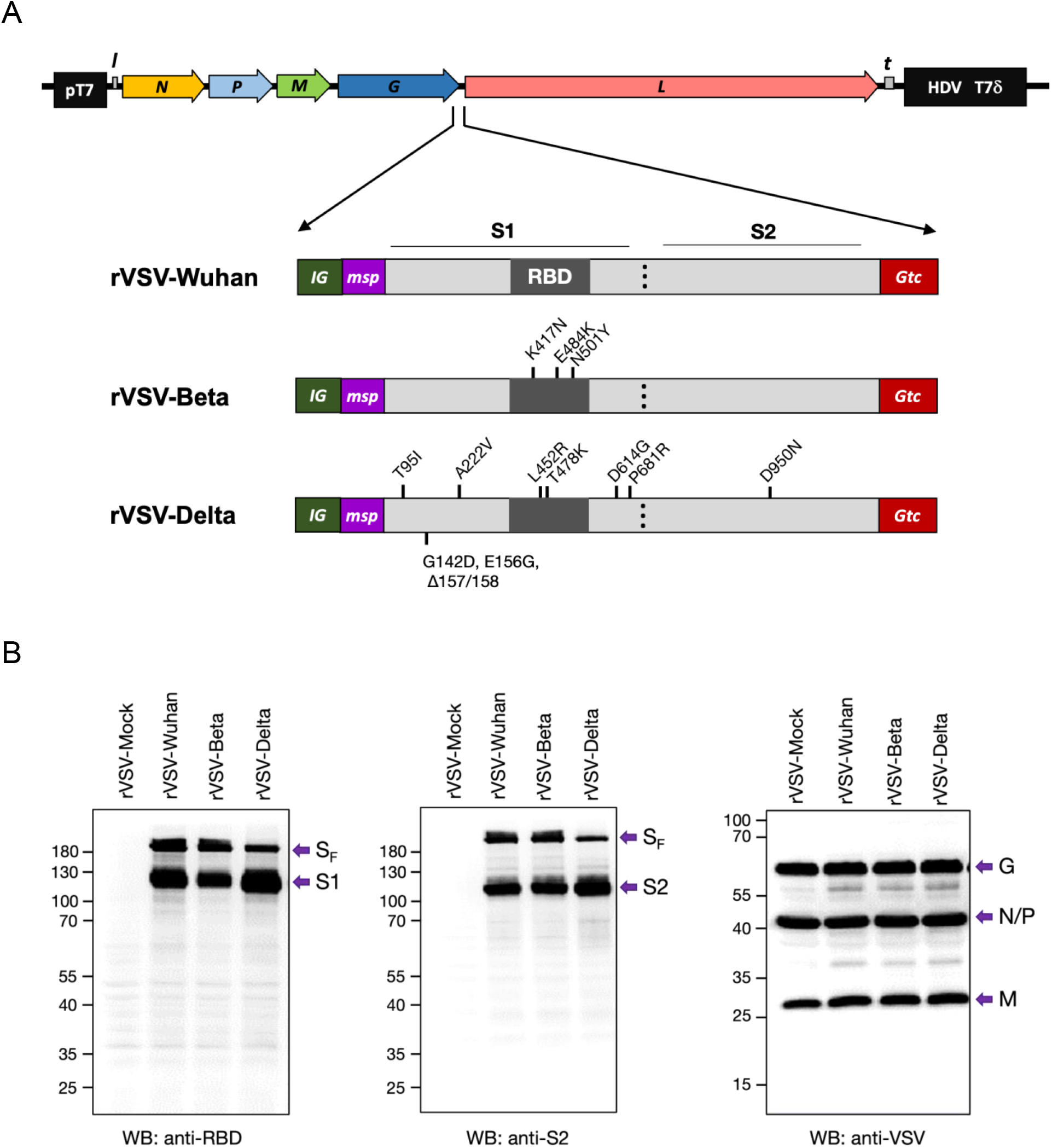
Construction of recombinant rVSV with S_F_ genes of SARS-CoV-2 Wuhan, Beta, and Delta. (A) Constructs of VSV-SARS-CoV-2 Wuhan (rVSV-Wuhan), Beta (rVSV-Beta), and Delta (rVSV-Delta). Codon-optimized full-length Spike protein gene (S_F_) with VSV intergenic sequences (IG), 21 amino acid honeybee melittin signal peptide (msp), and 49 amino acid VSV G protein transmembrane domain and cytoplasmic tail (Gtc) were inserted into the G and L gene junction of rVSV_Ind_. (B) Expression of SARS-CoV-2 Spike proteins in BHK-21 cells at 6 hr post-infection with rVSV-Mock (rVSV), rVSV-Wuhan, rVSV-Beta, and rVSV-Delta. pT7: Bacteriophage T7 promoter for DNA dependent RNA polymerase. N: VSV Nucleocapsid protein gene. P: VSV Phosphoprotein gene. M: VSV Matrix protein gene. G: VSV Glycoprotein gene. L: VSV Large protein, RNA dependent RNA polymerase gene. l: Leader region in the 3’
s-end of the VSV genome. t: Trailer region in the 5’-end of the VSV genome. HDV: Hepatitis delta virus ribozyme encoding sequences. T7δ: Bacteriophage T7 transcriptional terminator sequences. nt: nucleotides. aa: amino acids.

### Western Blot Analysis

We analyzed the expression of SARS-CoV-2 S proteins (S_F_, S1, and S2) in infected BHK-21 cells by Western blot analysis. Briefly, BHK-21 cells in 6-well plates were infected with recombinant rVSV-Wuhan, rVSV-Beta, and rVSV-Delta vaccines at a multiplicity of infection (MOI) of 6. At six hours post-infection, we lysed infected cells in 100 μl of lysis buffer (10 mM Tris-HCl, pH 7.4, 1% Nonidet P-40, 0.4% Na-deoxycholate, 10 mM Na_2_EDTA). Ten µg of the cell lysates were used for the Western blot analysis. SARS-CoV-2 S1 and S_F_ were detected by a rabbit polyclonal anti-SARS-CoV-2 Spike RBD antibody (Sino Biological, 40592-T62). S2 protein was detected by rabbit polyclonal anti-SARS-CoV-2 Spike S2 antibody (Sino Biological, 40590-T62). VSV proteins were detected using rabbit antiserum against lysed VSV Indiana (8).

### SARS-CoV-2 Viruses

SARS-CoV-2, Isolate USA-WA1/2020 (SARS-CoV-2_Wuhan_USAWA1_), NR-52281, contributed by the Centers for Disease Control and Prevention; SARS-CoV-2, Isolate hCoV-19/South Africa/KRISP-K005325/2020 (B.1.351; SARS-CoV-2_Beta_), NR-54009, contributed by Alex Sigal and Tulio de Oliveira; and SARS-CoV-2, Isolate hCoV-19/USA/PHC658/2021 (Lineage B.1.617.2; SARS-CoV-2_Delta_), contributed by Dr. Richard Webby and Dr. Anami Patel, were obtained through BEI Resources, NIAID, NIH. SARS-CoV-2, Omicron variant (BA.1; SARS-CoV-2_Omicron_) Isolate was obtained from the BC Centre for Disease Control (BCCDC) Public Health Laboratory, PHSA. For SARS-CoV-2 propagation, Vero E6 cells for SARS-CoV-2_Wuhan_USAWA1_, SARS-CoV-2_Beta_, and SARS-CoV-2_Delta_ or Vero cells for SARS-CoV-2_Omicron_, were infected with passage 1 virus for 3 days. The supernatant was harvested, cell debris pelleted and supernatant frozen. Serial dilutions of thawed virus-containing supernatants were used to infect Vero E6 or Vero cells to determine 50% tissue culture infectious dose (TCID50) via the Reed-Muench method (15).

### Animal Immunization

We prime-immunized 6 female BALB/c mice (10-12 weeks old; Jackson Laboratory) with 5×10^8^ PFU rVSV-Wuhan vaccine, -Beta vaccine, -Delta vaccine or 5×10^8^ PFU trivalent formulation (equal ratio rVSV-Wuhan vaccine, rVSV-Beta vaccine, rVSV-Delta vaccine). Immunizations were performed intramuscularly (i.m) on the hind leg. Three weeks after prime immunization, mice were administered monovalent or trivalent boost vaccines (5×10^8^ PFU i.m; **Figure 2**). Two weeks post boost, we collected blood via terminal cardiac puncture and isolated serum from the clotted blood after centrifugation. Spleens were collected and splenocytes isolated as previously described (16). Animal experiments were conducted in compliance with protocols (2020-084) approved by the Western University Animal Use Subcommittee.

**Figure 2:**
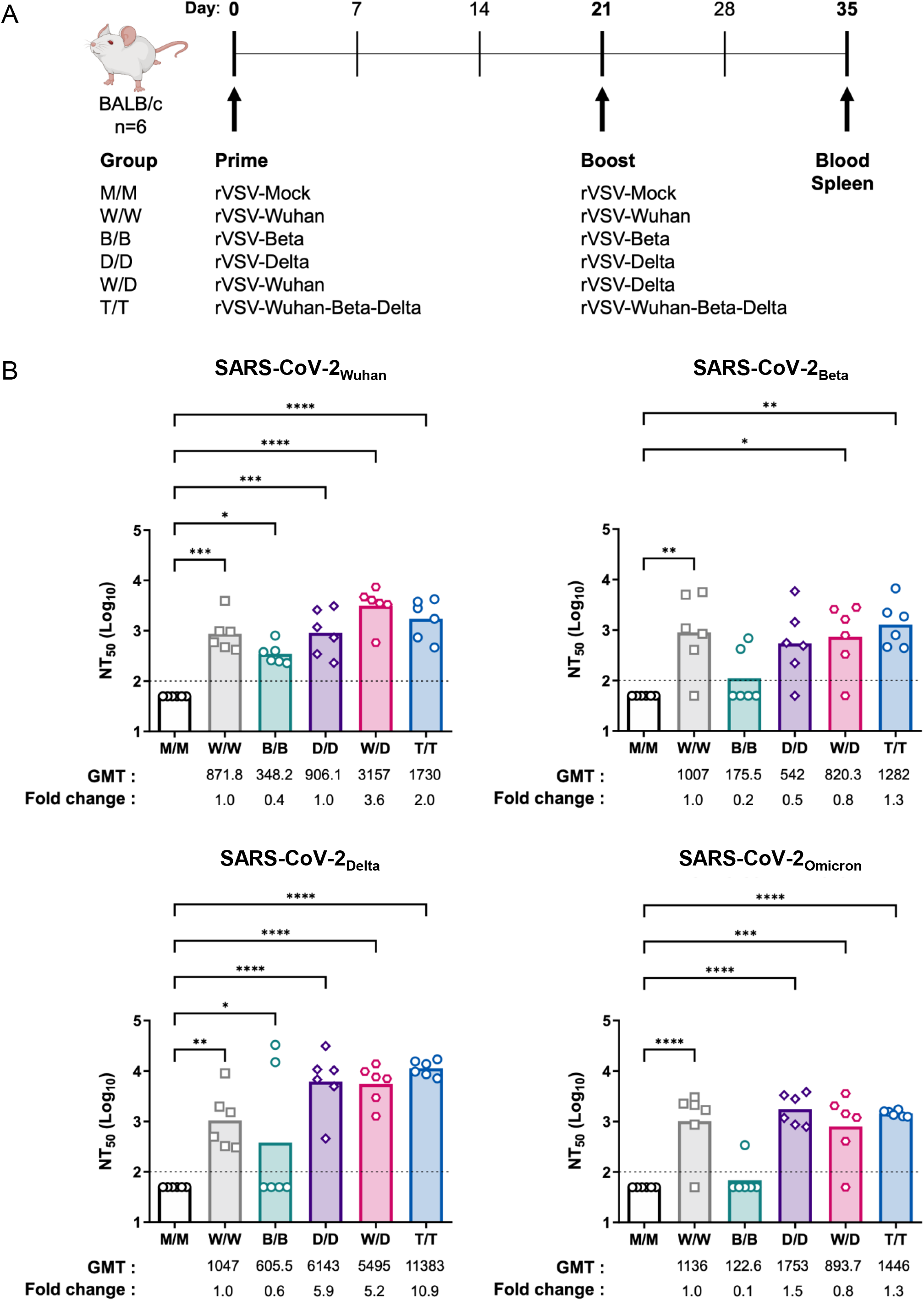
Immunization with monovalent and trivalent vaccines elicit a broad neutralizing antibody response. (A) Female BALB/c mice were prime immunized with 5×10^8^ PFU intramuscularly (i.m) of either rVSV-monovalent (rVSV-Mock [M], rVSV-Wuhan [W], rVSV-Beta [B], rVSV-Delta [D]) or trivalent (rVSV-Wuhan-Beta-Delta [T]) vaccine (day 0). On day 21 mice were administered a 5×10^8^ PFU homologous or heterologous boost i.m. Two weeks post-boost blood and spleen were collected for analysis of the immune response. (B) SARS-CoV-2 neutralization (SARS-CoV-2_Wuhan_USAWA1_, SARS-CoV-2_Beta_, SARS-CoV-2_Delta_, SARS-CoV-2_Omicron_) by immune sera two weeks post-boost was determined using replication-competent SARS-CoV-2 isolates in microneutralization assays. Graphed data are presented as arithmetic mean of log-transformed 50% neutralization titer (NT50). The horizontal dotted lines indicate the limit of detection. Each symbol denotes an individual animal; n=6; 2 independent experiments. Statistical significance was determined by one-way ANOVA with Bonferroni’s correction for multiple comparisons (*, p < 0.05; **, p<0.01; ***, p< 0.001; **** p<0.0001). Geometric mean NT_50_ (GMT) and the fold increases in GMT NT_50_ versus W/W (rVSV-Wuhan/Wuhan) are shown for each vaccine group.

### Microneutralization assay using replicating SARS-CoV-2 viruses

Microneutralization assays using replicating SARS-CoV-2 viruses were performed in VeroE6 cells for SARS-CoV-2_Wuhan_USAWA1_, SARS-CoV-2_Beta_, and SARS-CoV-2_Delta_; and Vero cells for SARS-CoV-2_Omicron_ as described previously (17). All procedures were performed in a BSL-3 facility at the Imaging Pathogens for Knowledge Translation (ImPaKT) facility, Western University following standard safety guidelines. Briefly, VeroE6 or Vero cells were seeded at 2×10^4^/well in 96-well cell culture plates in complete DMEM (10% FBS, 1x Pen/Strep) one day prior to infection. Heat inactivated serum samples (56°C for 30 min) were serially diluted (3-fold) in minimum essential media (MEM; Gibco) supplemented with 1x Glutamax (Gibco), 0.1% sodium bicarbonate (w/v, Gibco), 10 mM HEPES (Gibco), 1x penicillin/streptomycin (P/S; Wisent), 0.2% BSA (Sigma-Aldrich), and 2% fetal bovine serum (Wisent), starting at 1:100. Diluted sera were incubated with 250 TCID50/well of SARS-CoV-2 virus for one hour at RT, and 120 μl/well of the virus-sera mix was transferred to cells and incubated for 1 h at 37°C with 5% CO_2_. Virus was removed and replaced with 100 μl/well of the corresponding serum dilutions plus 100 μl/well of supplemented MEM. Plates were incubated for 72 h at 37°C with 5% CO_2_, then fixed with 10% formaldehyde solution (250 µl/well; Sigma-Aldrich) overnight at 4°C. Formaldehyde solution was removed, and cells were washed with PBS (pH 7.4), ready for nucleoprotein staining. Cells were permeabilized with 200 μl/well of PBS + 0.1% Triton X-100 (Sigma-Aldrich) for 15 min at RT, the permeabilization solution was removed, and plates were washed twice with 200 μl/well of PBS. Cells were blocked with PBS + 3% skim milk for 1 hour at RT before blocking buffer was removed and cells incubated for 1 hour with HRP-conjugated mouse anti-SARS nucleoprotein monoclonal antibody (1 µg/ml 1C7; Bioss Inc.) diluted in PBS with 1% skim milk (100 µl/well). Cells were washed twice with PBS, before addition of Sigmafast OPD (100 µl/well; Sigma-Aldrich) for 12 min at RT, followed by addition of 50 μl/well of 3 M HCl solution to stop reaction. Plates were read at 490 nm on a Cytation 5 microplate reader (BioTek). Analysis was performed using GraphPad Prism 9.3.1 software. Neutralization was calculated by subtraction of background (no virus) and then percentage of neutralization of sample wells relative to “virus only” control. To calculate the 50% neutralizing titer (NT_50_) a nonlinear regression curve fit analysis was performed on normalized data (top and bottom constraints set to 100% and 0% respectively).

### Intracellular IFN-γ staining

Splenocytes (2×10^6^ cells/condition) were stimulated for 5 hours ex vivo with 0.5 µM of MHCI restricted (S_535_-KNKCVNFNF; S_268_-GYLQPRTF; S_1052_-FPQSAPHGV) or 5 µM of MHCII restricted (S_61_-NVTWFHAIHVSGTNG; S_444_-KVGGNYNYLYRLFRK) synthetic peptides corresponding to BALB/c CD8+ and CD4+ T cell epitopes of SARS-CoV-2 Spike protein (18). IFN-γ production was assessed by intracellular flow cytometry as previously described (16). Briefly, negative controls were media alone and irrelevant peptide (MHCI restricted, lymphocytic choriomeningitis virus (LCMV) nucleoprotein NP_118–126_-RPQASGVYM). Brefeldin A (10 μg/mL, Sigma-Aldrich) was added to cells for the last 3 hours of T cell stimulation. Samples were washed in PBS and blocked with 2.4G2 hybridoma supernatant. Fluorescent conjugated anti-CD8α-Allophycocyanin (APC) mAb (clone 53-6.7, eBioscience) or APC-anti-CD4-APC mAb (clone GK1.5, eBioscience) were used for surface staining. Cells were fixed and permeabilized with Foxp3 / Transcription Factor Staining Buffer Set (eBioscience). Finally, cells were stained with a FITC–conjugated anti-IFN-γ mAb (clone XMG1.2, eBioscience). Frequency of IFN-γ+ T cells was assessed by on a BD FACS Canto II cytometer and analysed using FlowJo (Treestar).

### Statistics

Data were statistically analysed in GraphPad Prism 9.3.1. Normality of data was tested using a Shapiro-Wilk Test. Once confirmed, one-way ANOVA with Bonferroni post-tests vs. rVSV-Mock/Mock vaccine were undertaken to determine significance (p<0.05). Data was expressed on graphs as arithmetic mean with each data point representing a single mouse.

## Results

rVSV SARS-CoV-2 variant monovalent vaccines were generated by cloning Wuhan and variant spike protein genes (Beta and Delta) into rVSV vectors of serotype Indiana (8). The rVSV SARS-CoV-2 vaccine construct, described in (8), contains codon-optimized full-length SARS-CoV-2 Wuhan spike flanked by VSV intergenic sequences (IG) and 21 amino acid honeybee melittin signal peptide (msp) at the N-terminus and 49 amino acid VSV G protein transmembrane domain and cytoplasmic tail (Gtc) at the C-terminus, giving rise to rVSV-Wuhan vaccine. To generate variant vaccines, the SARS-CoV-2 Wuhan spike coding sequence was replaced with the SARS-CoV-2 Beta and Delta spike sequences generating the rVSV SARS-CoV-2 vaccines (rVSV-Beta vaccine, rVSV-Delta vaccine) utilized herein (**Figure 1A**). The expression of SARS-CoV-2 variant spike protein was confirmed via western blot for spike proteins (S1 and S2) and VSV proteins (**Figure 1B**). Highly purified vaccine virions containing variant spikes were generated for mouse vaccination (5×10^8^ PFU/mouse). Equal mix of the rVSV-Wuhan, rVSV-Beta and rVSV-Delta monovalent vaccines (1.67×10^8^ PFU/vaccine) produced a trivalent vaccine (total 5×10^8^ PFU/mouse) that was used in a homologous prime-boost regimen. Female BALB/c mice were prime immunized with 5×10^8^ PFU of the monovalent or trivalent vaccines via intramuscular injection and received a homologous or heterologous boost 21 days later (**Figure 2A**). Blood and spleen were collected 14 days post-boost (day 35) for detection of nAbs and cellular immunity, respectively (**Figure 2A**).

A live SARS-CoV-2 virus microneutralization assay detecting intracellular SARS-CoV-2 nucleocapsid protein (17), was used to test nAb efficacy of all the vaccine regimens against SARS-CoV-2_Wuhan_USAWA1_, SARS-CoV-2_Beta_, SARS-CoV-2_Delta_ and SARS-CoV-2_Omicron_. Prime-boost with the original rVSV-Wuhan monovalent vaccine (rVSV-Wuhan/Wuhan) induced an effective nAb response against matched SARS-CoV-2_Wuhan_USAWA1_. The remaining rVSV prime-boost combinations, rVSV-Beta/Beta, rVSV-Delta/Delta, rVSV-Wuhan/Delta and rVSV-Trivalent/Trivalent vaccines, also exhibited significant neutralization of SARS-CoV-2_Wuhan_USAWA1_, with rVSV-Wuhan/Delta and rVSV-Trivalent/Trivalent vaccines exhibiting superior neutralization with a 3.6-fold and 2-fold increase from rVSV-Wuhan/Wuhan immunization, respectively (**Figure 2B**). Serum antibodies elicited from rVSV-Beta/Beta vaccination were inefficient at neutralizing matched SARS-CoV-2_Beta_. Interestingly, at least 5 of 6 mice from the other vaccine regimens were able to cross-neutralize SARS-CoV-2_Beta_, with 100% of rVSV-Trivalent/Trivalent immunized mice generating a neutralizing antibody response. (**Figure 2B**). SARS-CoV-2_Delta_was neutralized effectively by all vaccine regimens except rVSV-Beta/Beta vaccination (**Figure 2B**). Serum from rVSV-Delta/Delta immunization exhibited a 5.9-fold increase in SARS-CoV-2_Delta_ neutralization versus rVSV-Wuhan/Wuhan immunization, with the rVSV-Wuhan/Delta vaccine regimen exhibiting a similar increase (5.2-fold). Of note, the rVSV-Trivalent/Trivalent vaccine regimen was most effective with a 10.9-fold increase in neutralization of SARS-CoV-2_Delta_. The most highly mutated variant to date, SARS-CoV-2_Omicron_, was neutralized by nAbs induced through prime-boost of rVSV-Wuhan/Wuhan vaccine, with heterologous boost (rVSV-Wuhan/Delta vaccines) being equally as effective. Prime-boost with rVSV-Delta/Delta vaccine was most effective with 1.5-fold increase in neutralizing titer, followed by a 1.3-fold increase with rVSV-Trivalent/Trivalent regimen (**Figure 2B**). Overall, serum from rVSV-Trivalent/Trivalent immunization was effective at generating nAb responses in 100% of mice for all SARS-CoV-2 viruses tested (SARS-CoV-2_Wuhan_USAWA1_, SARS-CoV-2_Beta_, SARS-CoV-2_Delta_ and SARS-CoV-2_Omicron_).

Using peptide epitopes characterised in SARS-CoV-2 infection of BALB/c mice (18) (**Supplementary Table 1**), we analyzed CD4+ and CD8+ T cell responses 14 days post-boost through ex vivo peptide stimulation of splenocytes and detection of IFN-γ expression (**Figure 3A**). All rVSV prime-boost vaccine regimens elicited a significant cognate CD8+ T cell response to the MHCI-restricted peptide S_535_ when compared to rVSV-Mock/Mock immunization (**Figure 3B**). The greatest CD8+ T cell response was seen in the rVSV-Wuhan/Wuhan and rVSV-Trivalent/Trivalent immunized mice. IFN-γ production was not detected 14 days post-boost for two additional MHCI-restricted peptides and two MHCII-restricted peptides (**Supplementary Figure 1; Supplementary Table 2**). Overall, this indicates that all vaccinated mice mounted a SARS-CoV-2 spike-specific CD8+ T cell response.

**Figure 3:**
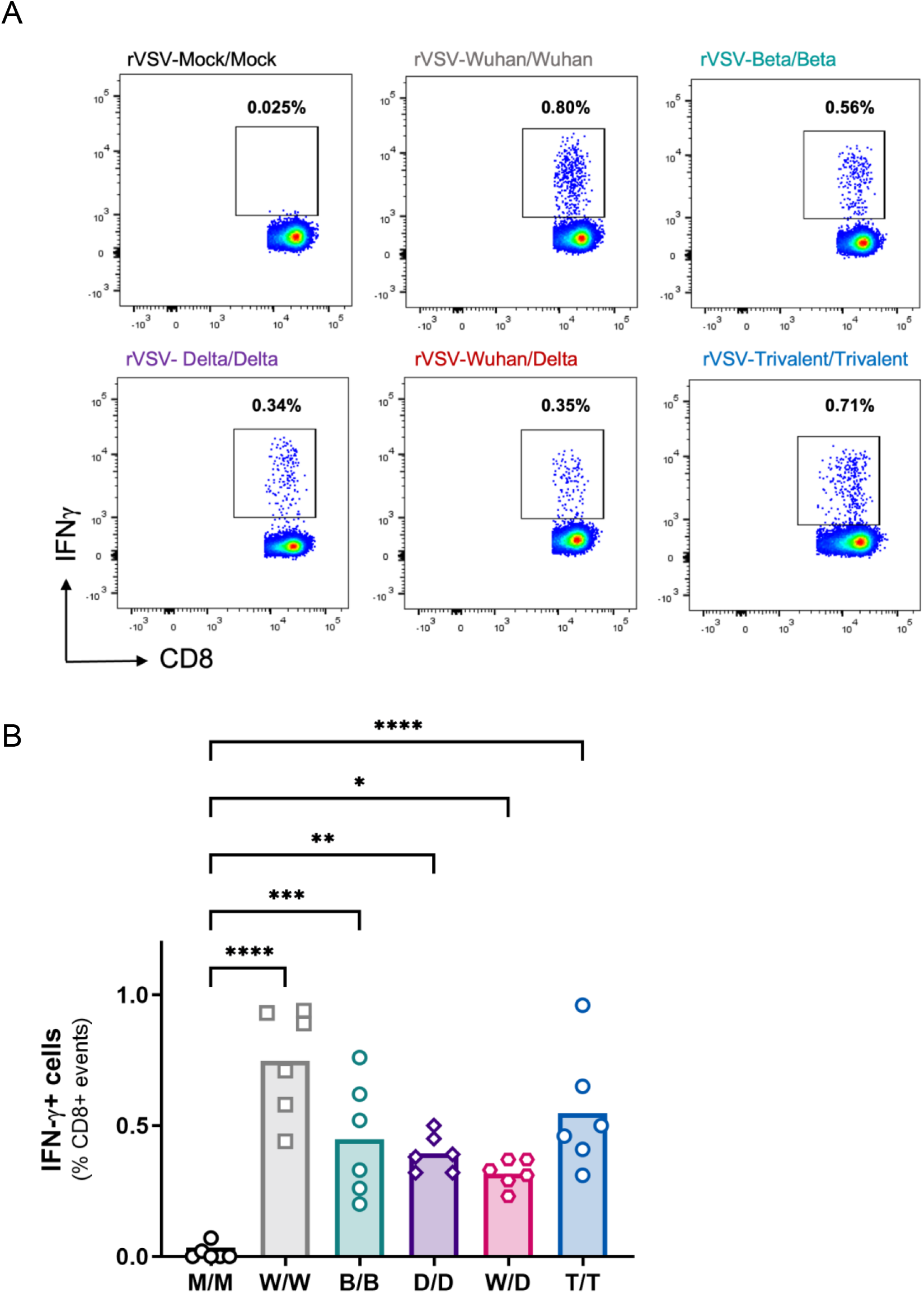
Monovalent and Trivalent vaccines generate a cognate CD8+ T cell response. Female BALB/c mice were prime immunized with 5×10^8^ PFU intramuscularly (i.m) of either rVSV-monovalent (rVSV-Mock, rVSV-Wuhan, rVSV-Beta, rVSV-Delta) or trivalent (rVSV-Wuhan-Beta-Delta) vaccine (day 0). On day 21 mice were administered a 5×10^8^ PFU homologous or heterologous boost i.m. Two weeks post-boost splenocytes were prepared and stimulated with SARS-CoV-2 specific epitope (S_535_ – KNKCVNFNF) ex vivo. IFN-γ production was determined following Brefeldin A treatment via intracellular cell staining and flow cytometry. (A) Representative dot plots of gated CD8+ T cells showing % IFN-γ+ cells of CD8+ events. (B) Pooled data from all animals for each vaccination group. Data represented arithmetic mean % IFN-γ+ cells of CD8+ events. Each symbol denotes an individual animal; n=6; 2 independent experiments. Statistical significance was determined by one-way ANOVA with Bonferroni’s correction for multiple comparisons (*, p < 0.05; **, p<0.01; ***, p< 0.001; **** p<0.0001). M/M: rVSV-Mock/Mock. W/W: rVSV-Wuhan/Wuhan. B/B: rVSV-Beta/Beta. D/D: rVSV-Delta/Delta. W/D: rVSV-Wuhan/Delta. T/T: rVSV-Trivalent/Trivalent.

## Discussion

Here we describe the first replication-competent rVSV vaccines containing variant SARS-CoV-2 spike genes. These SARS-CoV-2 variant vaccines used the rVSV platform previously validated as an effective prime boost regimen, containing the spike gene of Wuhan virus (8). In this study, effective nAb responses were elicited with all vaccine immunizations, as well as a spike-specific CD8+ T cell response, inducing both arms of the adaptive immune system. Homologous prime-boost of monovalent rVSV-Wuhan vaccine produced immune sera capable of neutralizing its matched SARS-CoV-2_Wuhan_USAWA1_, recapitulating our previous study, as well as all SARS-CoV-2 variant viruses tested, including the highly mutated SARS-CoV-2_Omicron_. The novel rVSV-Beta and rVSV-Delta vaccines generated nAbs against SARS-CoV-2_Wuhan_USAWA1_, however rVSV-Beta/Beta prime-boost immunization was ineffective at neutralizing matched SARS-CoV-2_Beta_ and more mutated variants. Other studies which tested monovalent Beta vaccines, in the Newcastle Disease Viral vector or mRNA vaccine platform, showed neutralization of SARS-CoV-2 B.1.351 (19,20). Expression of Beta spike was confirmed in our rVSV-Beta vaccine, however the lack of nAbs generated after immunization may be due to sequence differences between the immunizing rVSV-Beta vaccine spike, which lacked modifications (L18F, D80A and Del 242-244) located outside of the RBD, when compared to the SARS-CoV-2_Beta_ used in this study. Importantly, rVSV-Delta/Delta prime-boost immunization was effective at neutralizing matched SARS-CoV-2_Delta_ and superior to all vaccines against SARS-CoV-2_Omicron_, suggesting it induces a cross-protective immune response. Of note, we evaluated the effectiveness of a variant rVSV-Delta vaccine booster in animals which had received rVSV-Wuhan vaccine prime. Heterologous rVSV-Delta vaccine boost exhibited enhanced neutralization of SARS-CoV-2_Wuhan_USAWA1_ and SARS-CoV-2_Delta_ compared to homologous prime-boost with the original rVSV-Wuhan. Effectiveness of the rVSV-Wuhan/Delta prime-boost immunization regimen against SARS-CoV-2_Omicron_ was equivalent to homologous rVSV-Wuhan prime-boost immunization, indicating that the heterologous boost did not enhance nAb breadth against the highly modified Omicron variant. Our unique formulation of a rVSV-Trivalent vaccine, containing equal rVSV-Wuhan, rVSV-Beta and rVSV-Delta vaccines consistently produced a nAb response against all SARS-CoV-2 viruses tested, with 100% of mice generating nAbs. rVSV-Trivalent homologous prime-boost immunization exhibited superiority in neutralizing SARS-CoV-2_Delta_ and had similar neutralizing titers against SARS-CoV-2_Omicron_ compared to all vaccines (except rVSV-Beta/Beta immunization), suggesting the induction of a broad cross-neutralizing immune response.

Our rVSV-Trivalent vaccine was formulated to test whether the presence of multiple, varied spike sequences in a vaccine would induce cross-protective immune responses against unmatched and distinct emerging lineages of SARS-CoV-2. This approach has been utilized with multiple vaccine platforms against SARS-CoV-2 (19–21). Bivalent mRNA boosters developed combining Wuhan and Beta sequences, showed effective neutralization of SARS-CoV-2 B.1.612.7 and SARS-CoV-2 B.1.529 at day 29 post-boost (19), with a response detectable at day 181, albeit neutralizing titers were less against SARS-CoV-2 B.1.529 when compared to SARS-CoV-2 B.1.617.2 (19). Multivalent vaccines using the Newcastle virus as the vaccine vector have tested trivalent (Wuhan, Beta, Delta) and tetravalent (Wuhan, Beta, Delta, Gamma) formulas, with an increase to four different spike variants in the tetravalent exhibiting only a modest increase in nAb titer that did not translate to increased protection following challenge with SARS-CoV-2 Wuhan, Delta or Mu viruses (20). In all these studies, and ours, the contribution of each variant spike to the immune response is not clear. One study tested for the differential IgG detection of variant spikes with immune sera induced following monovalent and multivalent immunization of animals, showing induction of broad and consistent antibody cross-reactivity across all vaccine regimens (20), failing to decipher the contribution of each spike to the immune response. Our rVSV-Trivalent prime-boost vaccine regimen provided high, consistent, and broad neutralization of all variants tested, from SARS-CoV-2_Wuhan_USAWA1_ to the most recent variant of concern, SARS-CoV-2_Omicron_. Whether one spike sequence or a conserved immunodominant epitope is responsible for the cross-neutralization seen with multivalent vaccines is yet to be determined. We do know however, that our vaccines induced an effective CD8+ T cell response indicated by IFN-γ production stimulated by spike peptide (S_535_), conserved in all SARS-CoV-2 variants (18). This conserved epitope, located in the RBD of SARS-CoV-2, induced IFN-γ production in all monovalent and trivalent immunization groups. Additional peptides known to be immunodominant in BALB/c mice did not activate T cells ex vivo at two weeks post-boost, which may be because the peak of the T cell response one week post-boost had passed (18). Regardless, effective nAb and cognate CD8+ T cell responses were induced with each vaccination group, with future studies to assess how this translates into protection from SARS-CoV-2 challenge to be undertaken.

In our study, rVSV monovalent and trivalent vaccine regimens were effective in inducing a nAb response against SARS-CoV-2_Omicrom_, with cross-reactivity even occurring in the rVSV-Wuhan prime-boost immunization group. This is different from other studies where the BriLife® rVSV vaccine exhibited reduced effectiveness against SARS-CoV-2 B.1.529 versus the original SARS-CoV-2 virus, notably this vaccine had spontaneously acquired mutations evident in some SARS-CoV-2 variants (10). Moreover, the Newcastle-based monovalent Wuhan spike vaccine did not neutralize Omicron, but neutralization was achieved with trivalent and tetravalent formulations (20). The effectiveness of our vaccines against SARS-CoV-2 Omicron suggests inherent advantages of the rVSV vaccine platform. Our vaccines, unlike other SARS-CoV-2 rVSV vaccines (10,11), maintain the expression of VSV’s glycoprotein, generating a chimeric VSV-glycoprotein and SARS-CoV-2 spike vaccine (8). The presence of the VSV glycoprotein enhances replication of the rVSV vaccine in the recipient, lending itself to be more immunogenic than those that completely replace VSV glycoprotein. Our Wuhan and variant spike sequences also contained additions at the amino and carboxy terminus (msp and Gtc, respectively) that were shown to enhance spike expression, virion incorporation and immunogenicity (8), likely contributing to their broad efficacy.

We have utilized a validated rVSV-Wuhan vaccine platform to develop new rVSV vaccines to protect against SARS-CoV-2 variants that continue to emerge. The most effective vaccine regimens included rVSV-Wuhan/Delta heterologous prime-boost immunization, rVSV-Delta/Delta prime-boost immunization, and our unique rVSV-Trivalent/Trivalent prime-boost vaccine series. Further validation of whether a monovalent or multivalent approach is more effective for these vaccines, and determination of what spike sequences would be optimal in a multivalent vaccine are ongoing. The ability of these monovalent and trivalent rVSV variant vaccines to cross-neutralize the original SARS-CoV-2_Wuhan_USAWA1_ virus and those that continue to emerge, reveal their potential as effective booster vaccines for a population largely prime and boost-immunized with Wuhan spike-based mRNA vaccines.

## Acknowledgements

We would like to thank Richard Gibson and Dr. Amanda Hamilton from the Imaging Pathogens in Knowledge Translation (ImPaKT) Facility at Western University for their assistance during this study. We thank the veterinarians and animal care staff at the Health Sciences Animal Care Facility at Western University for their excellent husbandry of our experimental animals. Funding for this study was provided by CIHR of Canada and the Public Health Agency of Canada through a COVID-19 Immunity Taskforce grant (2020-VR2-173205) to RMT; a grant from CIHR (COV-440388) to SDB, RMT, EJA, SMMH, and CYK; a CoVaRR-Net Rapid Response Research Grant to SDB; and from Sumagen Canada to CYK.

---

**Supplementary Figure 1:**
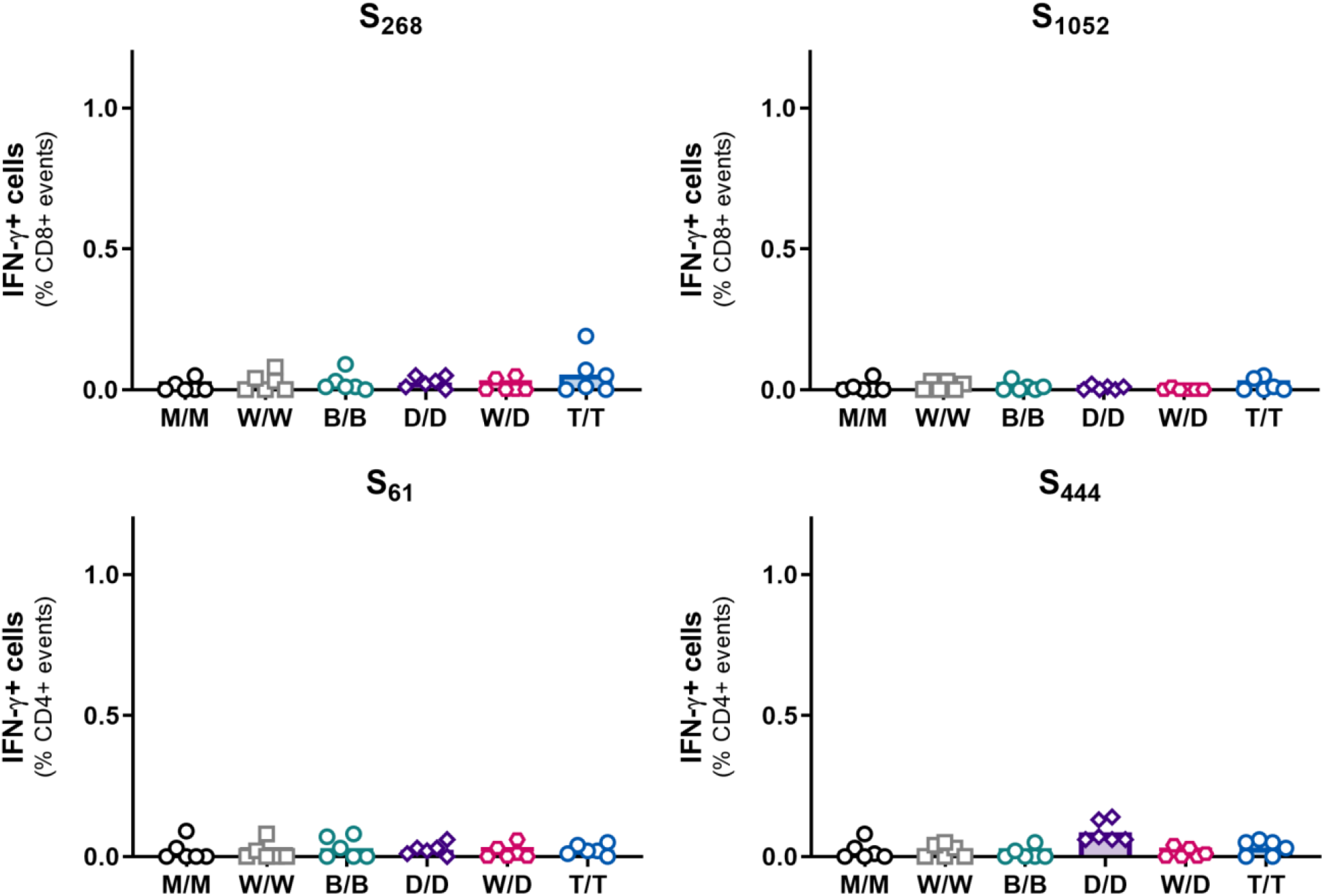
CD4+ and CD8+ T cell responses to spike-specific peptides. Female BALB/c mice were prime immunized with 5×10^8^ PFU intramuscularly (i.m) of either rVSV-monovalent (rVSV-Mock, rVSV-Wuhan, rVSV-Beta, rVSV-Delta) or trivalent (rVSV-Wuhan-Beta-Delta) vaccine (day 0). On day 21 mice were administered a 5×10^8^ PFU homologous or heterologous boost i.m. Two weeks post-boost, splenocytes were prepared and stimulated with SARS-CoV-2 specific epitopes ex vivo. CD8+ peptides included S_268_ (GYLQPRTF) and S_1052_ (FPQSAPHGV). CD4+ peptides included S_61_ (NVTWFHAIHVSGTNG) and S_444_ (KVGGNYNYLYRLFRK). IFN-γ production was determined following Brefeldin A treatment via intracellular cell staining and flow cytometry. Pooled data from all animals for each vaccination group. Data represented arithmetic mean % IFN-γ+ cells of CD8+ or CD4+ events. Each symbol denotes an individual animal; n=6; 2 independent experiments. Statistical significance was determined by one-way ANOVA with Bonferroni’s correction for multiple comparisons with no significant change seen from mock with all peptides. M/M: rVSV-Mock/Mock. W/W: rVSV-Wuhan/Wuhan. B/B: rVSV-Beta/Beta. D/D: rVSV-Delta/Delta. W/D: rVSV-Wuhan/Delta. T/T: rVSV-Trivalent/Trivalent.

**Supplementary Table 1.** Characteristics of SARS-CoV-2 Spike-specific T cell epitopes in BALB/c mice.

| Name | Sequence | Start | End | Size | Restriction |
| --- | --- | --- | --- | --- | --- |
| S268 | GYLQPRTF | 268 | 275 | 8 | MHCI (H2-D<sup>d</sup>) |
| S535 | KNKCVNFNF | 535 | 543 | 9 | MHCI (H2-K<sup>d</sup>) |
| S1052 | FPQSAPHGV | 1052 | 1060 | 9 | MHCI (H-2L<sup>d</sup>) |
| S61 | NVTWFHAIHVSGTNG | 61 | 75 | 15 | MHCII (I-E<sup>d</sup>) |
| S444* | KVGGNYNYLYRLFRK | 444 | 458 | 15 | MHCII (I-A<sup>d</sup>) |

\* Not conserved in SARS-CoV-2 B.1.617.2 (SARS-CoV-2<sub>Delta</sub>) and rVSV-Delta vaccine, which contain L452R.

